# The Ontogenetic Development of Hemispheric Lateralization During Face Processing: A Functional Magnetic Resonance Imaging Pilot Study in 7- to 9-Year-old Children

**DOI:** 10.1101/818310

**Authors:** Franziska E. Hildesheim, Isabell Debus, Roman Kessler, Ina Thome, Kristin M. Zimmermann, Olaf Steinsträter, Jens Sommer, Inge Kamp-Becker, Rudolf Stark, Andreas Jansen

## Abstract

Face processing is mediated by a distributed neural network commonly divided into a “core system” and an “extended system”. The core system consists of several, typically right-lateralized brain regions in the occipito-temporal cortex, including the occipital face area (OFA), the fusiform face area (FFA) and the posterior superior temporal sulcus (pSTS). It was recently proposed that the face processing network is initially bilateral and becomes right-specialized in the course of the development of reading abilities due to the competition between language-related regions in the left occipito-temporal cortex (e.g., the visual word form area) and the FFA for common neural resources.

The goal of the present pilot study was to prepare the basis for a larger follow-up study assessing the ontogenetic development of the lateralization of the face processing network. More specifically, we aimed on the one hand to establish a functional magnetic resonance imaging (fMRI) paradigm suitable for assessing activation in the core system of face processing in young children at the single subject level, and on the other hand to calculate the necessary group size for the planned follow-up study.

Twelve children aged 7-9 years, and ten adults were measured with a face localizer task that was specifically adapted for children. Our results showed that it is possible to localize the core system’s brain regions in children even at the single subject level. We further found a (albeit non-significant) trend for increased right-hemispheric lateralization of all three regions in adults compared to children, with the largest effect for the FFA (estimated effect size d=0.78, indicating medium to large effects). Using these results as basis for an informed power analysis, we estimated that an adequately powered (sensitivity 0.8) follow-up study testing developmental changes of FFA lateralization would require the inclusion of 18 children and 26 adults.

## 1 INTRODUCTION

Face processing is mediated by a distributed neural network. This network is, as first outlined in the Haxby model (Haxby et al., 2000), often divided into a “core system” and an “extended system” (Bernstein & Yovel, 2015). The core system consists of several bilateral brain regions in the occipito-temporal cortex. These regions include the fusiform face area (FFA) in the middle fusiform gyrus, the occipital face area (OFA) in the lateral inferior occipital gyrus and the posterior superior temporal sulcus (pSTS). The OFA has often been associated with the processing of single physical features of faces including the eyes, the mouth and the nose (Gschwind et al., 2012). The FFA is responsible for the analysis of invariant aspects of the face, as for example face identity (Rossion, 2015). The pSTS is involved in the processing of dynamic changeable facial features, for instance eye-gaze, mouth movements and facial expressions (Ishai et al., 2005). Beyond the core system, there are a number of additional (not face-specific) regions that contribute to face processing, e.g., the inferior frontal gyrus (IFG), the amygdala, the insula, and the orbitofrontal cortex (OFC). This extended system of face processing comes into play if additional information is extracted from faces, e.g., emotions, biographical information and attractiveness.

The neural face processing network is distributed across both hemispheres, but typically shows a right-hemispheric dominance in adults. This finding first originated from studies of patients with acquired prosopagnosia, i.e., the inability to recognize the identity of faces following brain damage. A large proportion of patients suffering from acquired prosopagnosia had lesions in the posterior right hemisphere (for an overview, see Bukowski et al., 2013). Although bilateral lesions often lead to more severe impairments than unilateral damage, unilateral-right damage is often sufficient to cause these impairments. Over the last 20 years, the face processing network has been extensively investigated in adults, in particular with functional magnetic resonance imaging (fMRI). Functional neuroimaging studies confirmed the right-hemispheric dominance of the face-processing network. The right hemisphere typically shows stronger response to face stimuli, both in terms of the spatial extent of the activation and the strength of activity (Frässle et al., 2016a; Frässle et al., 2016c; Bukowski et al., 2013; Willems et al., 2010; Hemond et al., 2007).

However, far less is known about its ontogenetic development. Some studies reported a pattern of increasing face-selectivity in the FFA (e.g., Haist et al., 2013; Peelen et al., 2009; Golarai et al., 2007; Aylward et al., 2005; Gathers et al., 2004). Nonetheless, it is still a matter of debate at what age this face selectivity develops (Golarai et al., 2007; Aylward et al., 2005; Gathers et al., 2004). Other studies reported a developmental shift from a more distributed activation pattern in children to a more focused activation pattern in adults (e.g., Scherf et al., 2007; Passarotti et al., 2003). Less research has been performed on the development of the OFA. A positive correlation between the intensity of right OFA activation and age was found (Joseph et al., 2011). This finding is in line with an earlier study that found lower intensity of face-preferential activation within the right-hemispheric OFA for children (6-10 years) compared to adolescents (11-14 years) and adults (Scherf et al., 2007). Findings on pSTS engagement during face processing in children are mixed. Some studies reported no (Joseph et al., 2011) or reduced (Scherf et al., 2007) pSTS recruitment in children. Other studies found no activation differences between children (of at least 7 years) and adults (Cohen Kadosh et al., 2011; Golarai et al., 2007). Yet, other studies even reported stronger pSTS recruitment in children compared to adults (Haist et al., 2013). Taken together, the empirical evidence on the functional neuroanatomy of the core face processing network in children is inconsistent, indicating large variability in terms of the localization of the brain regions of the core-system and its activation strength.

The present study focused on the development of hemispheric lateralization. It has been speculated that right-hemispheric lateralization of the core face processing network is emerging during development from childhood to adulthood. However, it remains unclear, at what age right-dominance emerges and which factors drive this specialization. Recent neuroimaging studies suggested that right-hemispheric specialization for face processing is initiated when children learn to read and further increases through adolescence (Behrmann & Plaut, 2015; Dundas et al., 2013; Dehaene & Cohen, 2011). The development of reading abilities, typically starting at the age of six, is neuroanatomically associated with the so-called visual word form area (VWFA). The VWFA is located in the left-hemispheric fusiform gyrus. It is considered to be an essential area for reading and is hypothesized to be involved in identifying words and letters from lower-level shape images, prior to association with phonology or semantics (Dehaene & Cohen, 2011; Price & Devlin, 2003). It is thought to be highly competitive with the FFA for common neural resources during childhood (Dundas et al., 2013; Cantlon et al., 2011). Both regions show similar positions in the fusiform gyrus, with a slightly more anterior location of the FFA compared to the VWFA (Dien, 2009). In order to optimize the connectivity between orthographical representations in the VWFA with typically already left-lateralized language areas, the VWFA gets gradually lateralized to the left hemisphere (Behrmann & Plaut, 2015; Dehaene & Cohen, 2011). It is hypothesized that due to competition between language-biased left-hemispheric VWFA specialization and face representation in the left hemisphere, face representation that was initially bilateral is driven to become right-specialized in the course of development (Behrmann & Plaut, 2015). Compared to the FFA, only few studies investigated developmental changes of OFA and pSTS lateralization during face processing. In analogy to the shift of FFA lateralization, OFA and pSTS lateralization are also expected to be subject to a developmental shift from a more bilateral activation to right-specialization during development, especially as brain regions closely interacting with each other benefit from being located close to one another to minimize signal propagation distance between those regions (Behrmann & Plaut, 2015).

Other theories explaining an emerging shift to the right hemisphere during face processing focus on differences in the face processing style of children and adults. The right hemisphere is believed to be involved in a holistic processing of faces, whereas the left hemisphere is more specialized in the processing of single features (Meng et al., 2012; Rhodes et al., 1993; Hillger & Koenig, 1991). It is suggested that adults encode faces using a holistic strategy based on the configural information of the face, i.e., the spatial relations among the different facial features (Dundas et al., 2013; Aylward et al., 2005; Schwarzer, 2000). Children younger than ten years tend to encode faces using an analytic strategy by analyzing distinctive facial features (Schwarzer, 2000). This analytic strategy would suggest a stronger recruitment of the left hemisphere during face processing in children compared to adults (Meng et al., 2012).

The present study aimed at setting up an imaging paradigm to investigate the ontogenetic development of the core system of face processing, in particular the hemispheric lateralization of the FFA. Its first goal was to establish an fMRI task suitable for assessing activation in the core system of face processing in young children (aged 7-9 years). The second goal was to compare both activation strength and hemispheric lateralization of the brain regions in the core system between children and adults. We hypothesized that children would show, due to still developing neural specialization, reduced activity in all brain regions of the core system (i.e., bilateral OFA, FFA and pSTS) and reduced right-hemispheric lateralization. Since we did not have well-founded estimates for the expected effect size, the second part of the study had pilot character. We aimed to use these results as basis for an informed power analysis yielding the necessary group size for a larger follow-up study that assesses differences in the neural network of face processing between children and adults.

## 2 MATERIALS AND METHODS

### 2.1 Subjects

Participants were recruited through distribution of flyers and bulletins in public places and advertisement through the student mailing list of the Philipps-University of Marburg, Germany. Ten adults (3 females, 7 males; 24-45 years; mean age 32.1 ± 6.1 years) and 12 children were initially recruited for the study. Three children were excluded from the final analysis. One child aborted the measurements prematurely due to anxiety. The other two children were excluded due to high motion during the scanning session (see results). The final children sample therefore comprised nine children (2 females, 7 males), aged 7 to 9 years (9.0 ± 0.7 years).

All participants had normal or corrected-to-normal vision and had no history of psychiatric or neurological disorders. Self-reported right-handedness was used as selection criterion during the recruitment process. To ensure right-handedness, subjects were additionally asked to complete the Edinburgh Handedness Inventory Questionnaire (Oldfield, 1971). According to this questionnaire, all children were right-handed with a mean laterality quotient (LQ) of +89.8. In the adult sample 9 out of 10 subjects were right-handed with a mean LQ of +88.3. One adult subject turned out to be left-handed with a LQ of −33.3. We included this subject for the first analysis in which we assessed in how many subjects it was possible to localize the core system’s brain regions on the individual subject level. We excluded this subject from the second analysis in which we compared adults and children, since it is known that activation strength and hemispheric lateralization can be influenced by handedness (Frässle et al., 2016a; Bukowski et al., 2013; Willems et al., 2010).

As a variety of studies have revealed that autistic traits or a very low empathy can result in a distinct pattern of face processing (Jospe et al., 2018; Naumann et al., 2018), participants were asked to complete the Autism Spectrum Quotient Questionnaire (AQ; Baron-Cohen et al., 2001) and the Empathy Quotient Questionnaire (EQ; child-version: Empathy-Systemizing-Quotient Questionnaire/EQ-SQ-Child; Baron-Cohen & Wheelwright, 2004). The evaluation of EQ and AQ showed that all children scored in normal range. Among the adults, two of the subjects achieved a below-average EQ score and one of the subjects was above-average in the AQ score. However, as these participants did not show significantly different activation patterns in the core network, they were included in all analyses. To assess overall cognitive abilities in children, the brief version of the standardized Wechsler Nonverbal Scale of Ability Test (Wechsler & Naglieri, 2006) was applied. As none of the children displayed major cognitive deficits, all 9 children subjects were included in further analyses.

All subjects provided written informed consent after they were apprised in detail about the experimental set up and the study procedure. The study was approved by the local ethics committee of the Department of Psychology of the Justus-Liebig University in Giessen, Germany (reference number 2018-0024). In case of minor subjects, their parents provided informed consent. After study participation, adult subjects received an allowance of 10 Euros and children could choose between different toys.

### 2.2 MRI investigation of children

The children were invited to visit the MR scanner facilities a couple of days in advance to the scanning session to familiarize them with the MR scanner environment, to reduce anxiety and discomfort and to minimize occurrence of strong motion artifacts. The individual training session started with a chair circle, including the participating child, one or both of its parents, two instructors and a professional radiographer, who conducted the actual scanning session. A playful theoretical introduction gave insights into the experimental procedure and the fMRI method. The coloring book Paula in der Röhre (Bayer HealthCare, 2008) was used for explaining the procedure. In this booklet the fMRI technique and aspects to consider when lying in the MR scanner are taught in a child-friendly narrative. The book was sent to the child prior to the training session with the appeal to take a look at it. In the training session, the story of the booklet was discussed with the child and possible questions were answered. Children were made aware of the extreme importance of lying still during the scanning session, by making use of the comparison between motion artifacts and blurry photographs (Wilke et al., 2018). In addition, children were shown some exemplary images of face and house stimuli from the fMRI paradigm to give them an impression of the experimental task.

The fMRI study was embedded in a child-oriented setting, putting the whole experiment in a frame story. The child was told to imagine being an astronaut who is flying in a rocket (i.e., the MR scanner). To motivate the child, a cuddly toy was brought into the story, which accompanies the child as a co-astronaut during the whole training and actual scanning session. By using the notion of the narrow interior of the rocket, the child should lose discomfort induced by the tightness inside the MR scanner. Intense background noise was explained as noise produced by the rocket when speeding up. The importance of lying still inside the MR scanner was further underlined by the story as resting still is crucial for a smooth steering of the rocket. To bring the stimuli of the fMRI paradigm into the frame story, children were told that on their journey through the universe they encounter people living in their houses on different planets, showing different reactions when seeing the rocket passing.

After introducing the child to the frame story, it had the possibility to inspect the scanning room and the MR scanner. Together with the radiographer, the child could first view the scanner from the outside. Afterwards, it was invited to lie in the scanner to get a feeling for the tightness of the tube. The head coil was also mounted for test purposes, being explained as the helmet of the astronaut. On the day of the scanning session, children were reminded of the frame story and of things to consider when lying inside the scanner.

### 2.3 Experimental paradigm

For MRI data acquisition, participants laid in supine position in the scanner with their head first. Light inside and outside the scanner was switched off to strengthen the children’s feeling of being situated in a spacecraft in the universe. A response box with one button was fixated on the right thigh of the subjects for conducting a button-press task during the fMRI paradigm. To prevent motion artifacts, soft foam rubber pads were used for head fixation. Stimuli were presented via an MRI-compatible LCD screen that was positioned behind the MR scanner. Subjects viewed the paradigm through a 45° tilted mirror which was fixated at the head coil. All stimuli were presented using the software package Presentation (version 20.2, Neurobehavioral Systems, San Francisco, California, USA).

The face processing network was investigated using a face localizer paradigm in which subjects viewed either gray-scale faces with neutral, sad or fearful expressions in the activation condition or houses in the control condition in a blocked design (see Fig. 1 for details).

**Figure 1:**
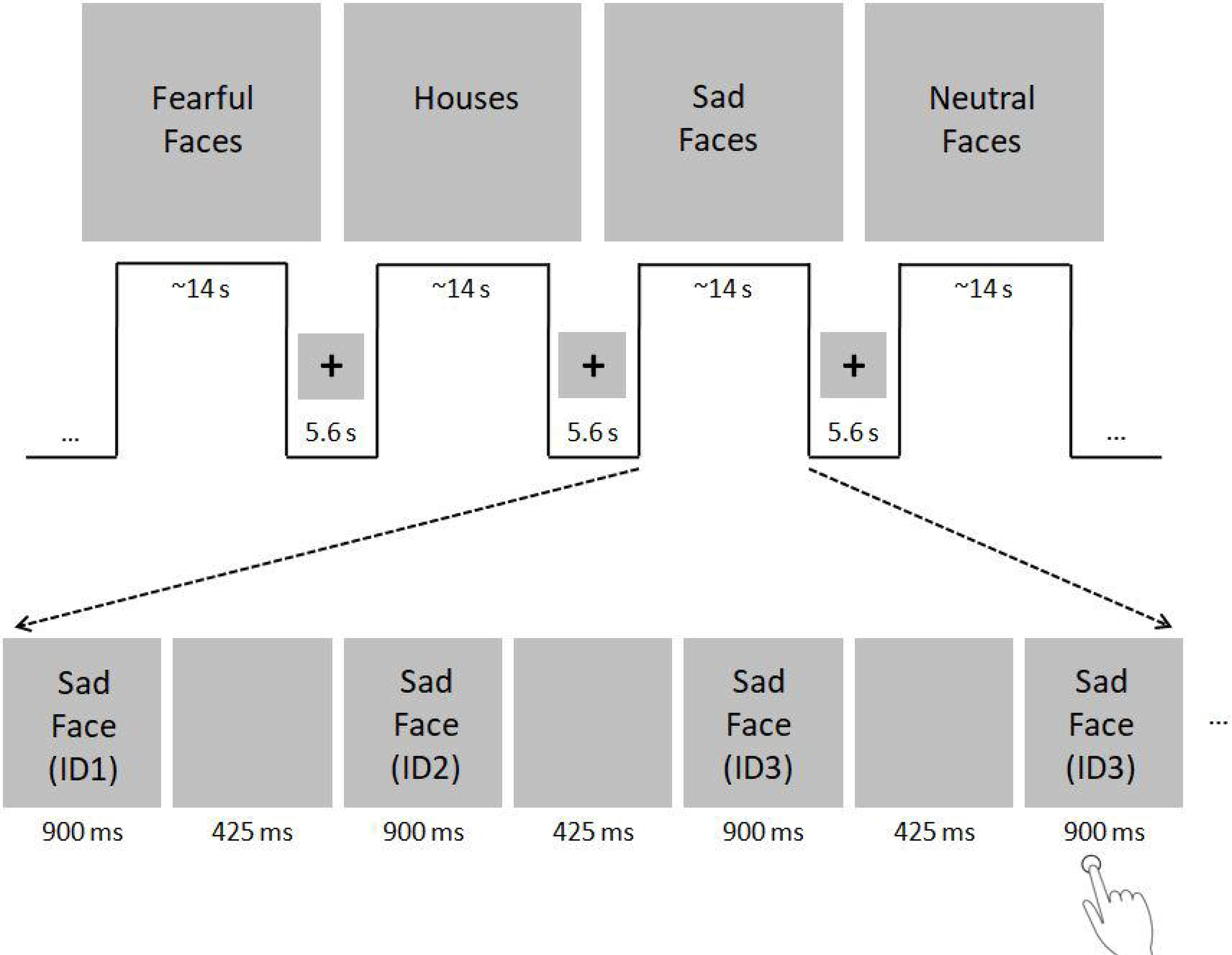
fMRI paradigm. Subjects viewed either gray-scale faces with neutral, sad or fearful faces in the activation condition and houses in the control condition in a blocked design. Face stimuli were selected from the Karolinska Directed Emotional Faces (KDEF) dataset (http://www.emotionlab.se/resources/kdef; Lundqvist et al., 1998). House stimuli were selected from the internet. The paradigm consisted of two sessions, including 16 blocks each (4 blocks with neutral, sad and fearful faces, respectively; 4 house blocks). The sessions were divided by a short break of 20 s. Each block included 11 stimuli that were presented for 900 ms with an inter-stimulus interval of 425 ms. Blocks lasted ~14 s each and were separated with blank periods (duration: 5.6 s) in which only a centered fixation cross was shown. The order of blocks remained the same across all subjects, whereas the order of images in each block was pseudo-randomized. To ensure attention, subjects were asked to indicate via button press with the right index finger when a stimulus was shown twice consecutively. Within one block, either two or three stimulus-pairs arose, which sum up to 40 target events in the whole fMRI paradigm. The total duration of the fMRI paradigm was ~11 minutes^4^.

### 2.4 MRI data acquisition

Subjects were scanned on a 3-Tesla MR scanner (Siemens Prisma 3-Tesla Magnetom) at the Bender Institute of Neuroimaging (BION) at the Justus-Liebig University of Giessen, Germany. All MRI data were acquired using a 64-channel head matrix receive coil.

First, a high-resolution anatomical image was acquired using a T1-weighted magnetization prepared rapid gradient echo (MPRAGE) sequence. The following parameters were applied: acquisition time (TA) 4:29 min, repetition time (TR) 1580 ms, echo time (TE) 2.30 ms, field of view (FOV) 240 mm, 176 slices, slice thickness (ST) 0.94 mm, resolution 0.9 × 0.9 × 0.9 mm, phase encoding direction (PE) anterior ≫ posterior, distance factor (DF) 50%, flip angle 8°, bandwidth 200 Hz/Px, sagittal ascending acquisition.

Second, functional images were collected using a T2*-weighted gradient echo-planar imaging (EPI) sequence sensitive to Blood Oxygen Level Dependent (BOLD) contrast. The following parameters were used: TA 11:14 min, TR 1780 ms, TE 36 ms, FOV 256 mm, 20 slices per slab, ST 2.4 mm, resolution 2.0 × 2.0 × 2.4 mm, PE anterior ≫ posterior, DF 20%, flip angle 70°, bandwidth 1396 Hz/Px, ascending acquisition. We did not measure the whole brain, but only a slab (Fig. 2). Reducing the coverage allows reduction of the voxel size and therefore an increased spatial resolution. The measurement of a slab of the brain is believed to facilitate the measurement of small regions (e.g., amygdalae; Morawetz et al., 2008). The slab was manually orientated, using the structural T1-weighted image. We aimed to cover on the one hand all three regions of the core face processing network, i.e., bilateral OFA, FFA and pSTS. On the other hand, we also aimed, as part of a related project, to measure activity in parts of the extended system, in particular the amygdala, insula, cingulate gyrus and inferior frontal gyrus. These brain regions are known to play an essential role in emotion processing across development.

**Figure 2:**
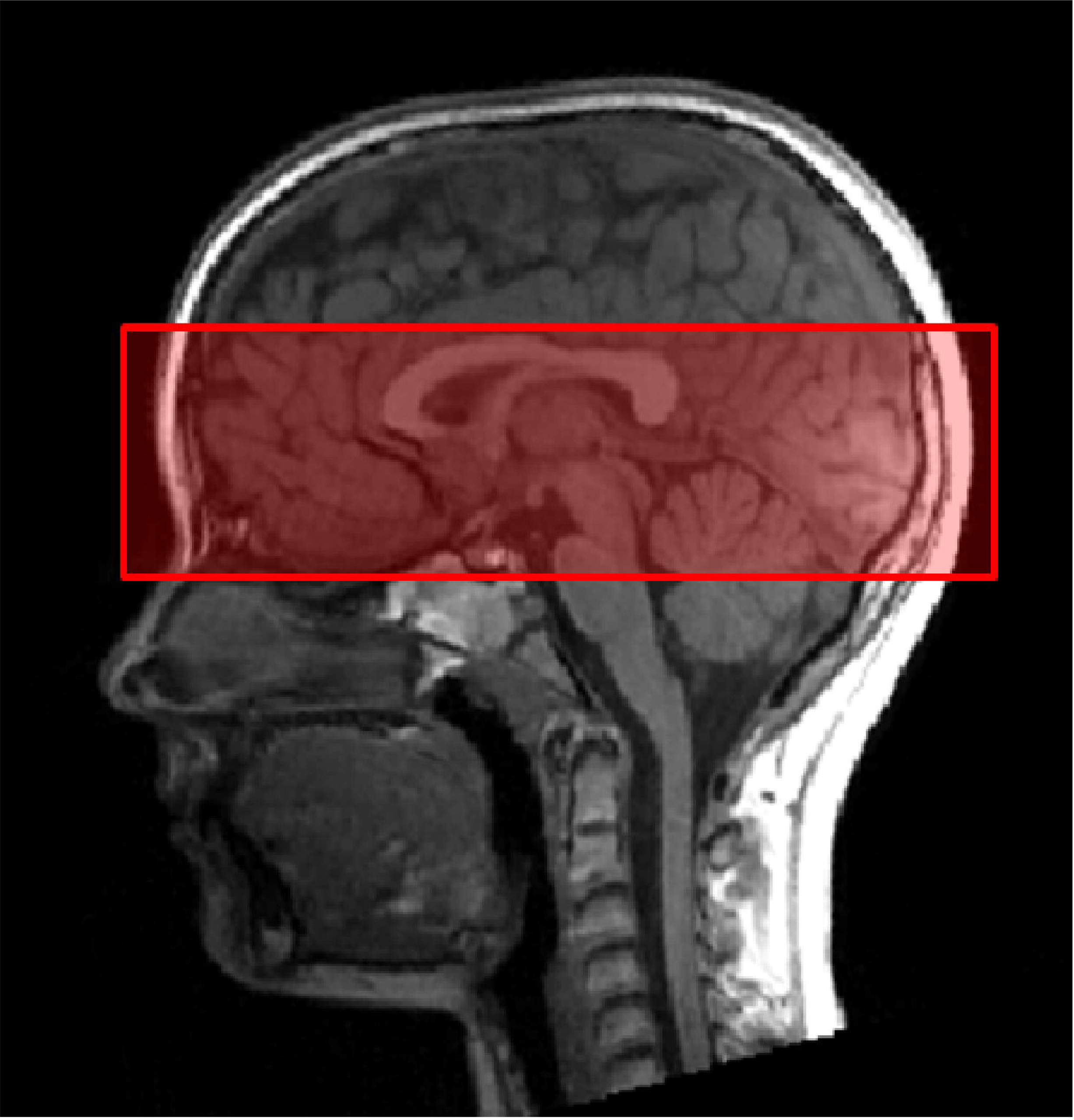
Slab orientation covering bilateral OFA, FFA and pSTS. The slab was oriented at the lowest part of the occipital pole and the lowest part of the prefrontal cortex (PFC), using the middle view of the structural image.

### 2.5 MRI data analysis

MRI data were analyzed using Statistical Parametric Mapping (SPM12, version 7219, Wellcome Trust Centre for Neuroimaging, London, UK), based on MATLAB (version 9.1, R2016b).

Preprocessing: To control for head movements, functional images from both sessions were realigned to the mean image. Realigned images were coregistered with the high-resolution anatomical image and then spatially normalized into the Montreal Neurological Institute (MNI) standard space using the unified segmentation-normalization of the anatomical image. Normalized functional images were spatially smoothed using an isotropic 6 mm full width at half maximum Gaussian kernel.

Statistical analysis was performed in a two-level, mixed-effects procedure. At the individual subject level, voxel-wise BOLD activity was modeled by a General Linear Model (GLM). Each condition of the face-localizer paradigm, i.e., neutral, sad and fearful faces, respectively, as well as houses, was modeled as a block regressor. This regressor was convolved with the hemodynamic response function implemented in SPM12. The regressors for the two sessions were entered into two separate sessions in one GLM (i.e., were not concatenated). In addition, the six realignment parameters of each session were included in the GLM design matrix as nuisance regressors to control for movement-related artifacts not accounted for by the realignment during preprocessing. A high-pass filter (cut-off frequency: 1/128 Hz) was used to account for low-frequency noise. Individual BOLD activity related to face processing was identified by a contrast comparing faces (irrespective of emotional content) against houses, averaged across sessions (i.e., using the contrast weights −3 1 1 1 0 0 0 0 0 0 −3 1 1 1 0 0 0 0 0 0). In the following, we will refer to this contrast as “faces > houses”. To assess brain activation at the group level, the “faces > houses” contrast images were entered separately for children and adults into one-sample t-tests. Anatomical localization of the activated brain regions was achieved using the WFU-Pickatlas (Maldjian et al., 2003).

Quality control: First, a motion analysis was performed to rule out that potential activation differences between children and adults were caused by unequal motion artifacts. The motion analysis was performed by MotionEstimator, developed by one of the authors (R.K.). MotionEstimator calculates location differences between two scans (see https://github.com/kesslerr/motionEstimator for details). As cut-off criterion for exclusion from further analysis, we chose a mean scan-to-scan motion exceeding 0.35 mm (Power, Schlaggar & Petersen, 2015). Second, further quality control was conducted using the software package MRIQC (Magnetic Resonance Imaging Quality Control; OpenNeuro.org, https://mriqc.readthedocs.io/en/stable/). MRIQC assesses both structural T1-weighted MR images and BOLD-images of the brain by calculating a set of quality measures from each image (Esteban et al., 2017). MRIQC uses 14 Image Quality Metrics (IQMs) that characterize each image in 56 features. The tool also includes a visual reporting system in order to manually investigate potential quality issues in single subjects.

### 2.6 Analysis strategy

First aim of the study was to assess whether it is possible to detect brain activity in the core system of face processing in children at the single subject level. We proceeded in two steps. First, we analyzed the group activation pattern for the contrast “faces > houses” separately for adults and children using one-sample t-tests. Second, we analyzed the individual activation pattern.

In these activation patterns, we determined whether brain activity could be found in the left OFA, right OFA, left FFA, right FFA, left pSTS and right pSTS. The brain activation patterns were first thresholded at a conservative threshold of p < 0.05, corrected for multiple comparisons at the whole brain level (family-wise error, FWE, corrected). If brain activity was not found in all regions of the core system at this threshold, the p-value was subsequently lowered to more liberal thresholds (p < 0.001 and p < 0.05, respectively, uncorrected for multiple comparisons) (see Schuster et al., 2017 for an extensive discussion of this procedure).

To assess whether or not a specific activation can be attributed to the core system of face processing, we created three anatomically defined regions of interest (ROIs) including bilateral OFA, FFA and pSTS, respectively, using the WFU-Pickatlas (Maldjian et al., 2003). OFA-ROI masks were created choosing the inferior occipital gyrus in the brain atlas IBASPM116 (as implemented in the WFU-Pickatlas). FFA-ROI masks were built choosing the fusiform gyrus. pSTS-ROI masks were created choosing the superior and middle temporal gyrus. Activation clusters that appeared inside one of the ROI masks were considered as potential candidates of core system brain activity. To verify the correct anatomical localization, both the anatomical localization on the canonical single-subject T1-image (implemented in SPM12) and the positions of the activated brain regions in the occipito-temporal lobe relative to each other were used. This identification procedure was performed by four individual raters (authors F.E.H., I.D., R.K., K.M.Z.) separately to maximize accuracy and minimize error-proneness due to intra-rater differences.

Second aim of the study was to compare activation strength and hemispheric lateralization of brain regions in the core system of face processing between adults and children.

Activation strength: We decided against applying the standard approach for a group analysis, i.e., assessing voxel-wise differences in normalized functional images between both groups using a two-sample t-test. Since the brains of adults and children largely differ in size and shape, the normalization procedure might have introduced systematic differences between these groups. Instead, we created for each subject individual spherical masks (radius 6 mm) centered at the corresponding local maximum of the six brain regions of the core system. Activation strength was calculated as the mean value of all voxels inside the respective mask for the weighted ß-image (contrast “faces > houses”)1. Activation differences between both groups were assessed with Welch-tests in R (www.r-project.org/">).

Lateralization: The degree of regional face-sensitive hemispheric lateralization was assessed by a lateralization index (LI) (Jansen et al., 2006). The LI is given by the following expression

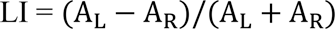

where AL and AR refer to values of fMRI-measured activity for homologous ROIs within the left (L) and right (R) hemisphere. The LI yields values between 1 and −1. In the present study, an LI > 0.20 was considered to represent left-hemispheric dominance and an LI < −0.20 right-hemispheric dominance. An LI between −0.20 and 0.20 was denoted as bilateral.

For calculation of the LI, we used the bootstrap procedure implemented in the LI tool-box extension (Wilke & Schmithorst, 2006; Wilke & Lidzba, 2007). This method takes 100 bootstrapped samples (resampling ratio k = 0.25) for the ROIs in the left and right hemisphere for 20 equally sized thresholds ranging from 0 to the maximum t-value. This results in 10,000 possible LI combinations, from which only the central 50% are kept to exclude statistical outliers. For each subject, a representative LI is then calculated by weighting these central 50% LIs with their respective threshold. Using this procedure, the LI was computed for face-sensitive activation in the OFA, FFA and pSTS. To specify the ROIs for LI calculation, individual masks for each subject’s bilateral OFA, FFA and pSTS were generated using the WFU-Pickatlas. Individual masks were created as spheres of 10 mm radius around the previously identified MNI-coordinates of each ROI. Lateralization differences between both groups were assessed with Welch-tests.

Power analysis: Since we did not have empirical estimates for the expected effect size of activation and lateralization differences between children and adults, the present results were intended to build the basis for an informed power analysis yielding the necessary group size for a larger follow-up study. The power analysis was performed using the software G*Power (version 3.1; Faul et al., 2009). The sample effects of different hypotheses detected in the present study were used as an estimate of the population value of the effect to be detected in consecutive studies (Anderson et al., 2017). For calculation, statistical power was set to 0.8 (80%) and the alpha error probability was set to α=0.05 (p < 0.05). Necessary sample sizes were calculated for both activity and lateralization measures. Effect sizes were calculated from the particular mean effect of the children sample, the mean of the adult sample and the standard deviation of each group. We used an unbalanced adults/children allocation ratio of 1.4. To achieve the same power, unbalanced designs have to include more subjects than balanced study designs. We nevertheless decided to perform the power analysis for this design, since it allowed us to reduce the number of children (difficult to recruit) at the cost of including overall more adult subjects (easy to recruit).

## 3 RESULTS

### 3.1 Motion analysis

Motion analysis was performed for all 10 adults and 11 children. None of the adult subjects showed a mean scan-to-scan motion exceeding the defined cut-off score of > 0.35 mm (Fig. 3 bottom). They showed an averaged mean scan-to-scan motion of 0.11 mm ± 0.05 mm in the first run and 0.12 mm ± 0.06 mm in the second run. However, two children showed a mean scan-to-scan motion exceeding > 0.35 mm in the second run (C03, C10; Fig. 3 top). They were therefore excluded from further analyses. Average mean scan-to-scan motion of the remaining 9 children was 0.09 mm ± 0.03 mm in the first run and 0.11 mm ± 0.05 mm in the second run. The overall motion of the included subjects was therefore comparable between children and adults. This rules out that potential activation differences between children and adults are caused by unequal motion artifacts.

**Figure 3:**
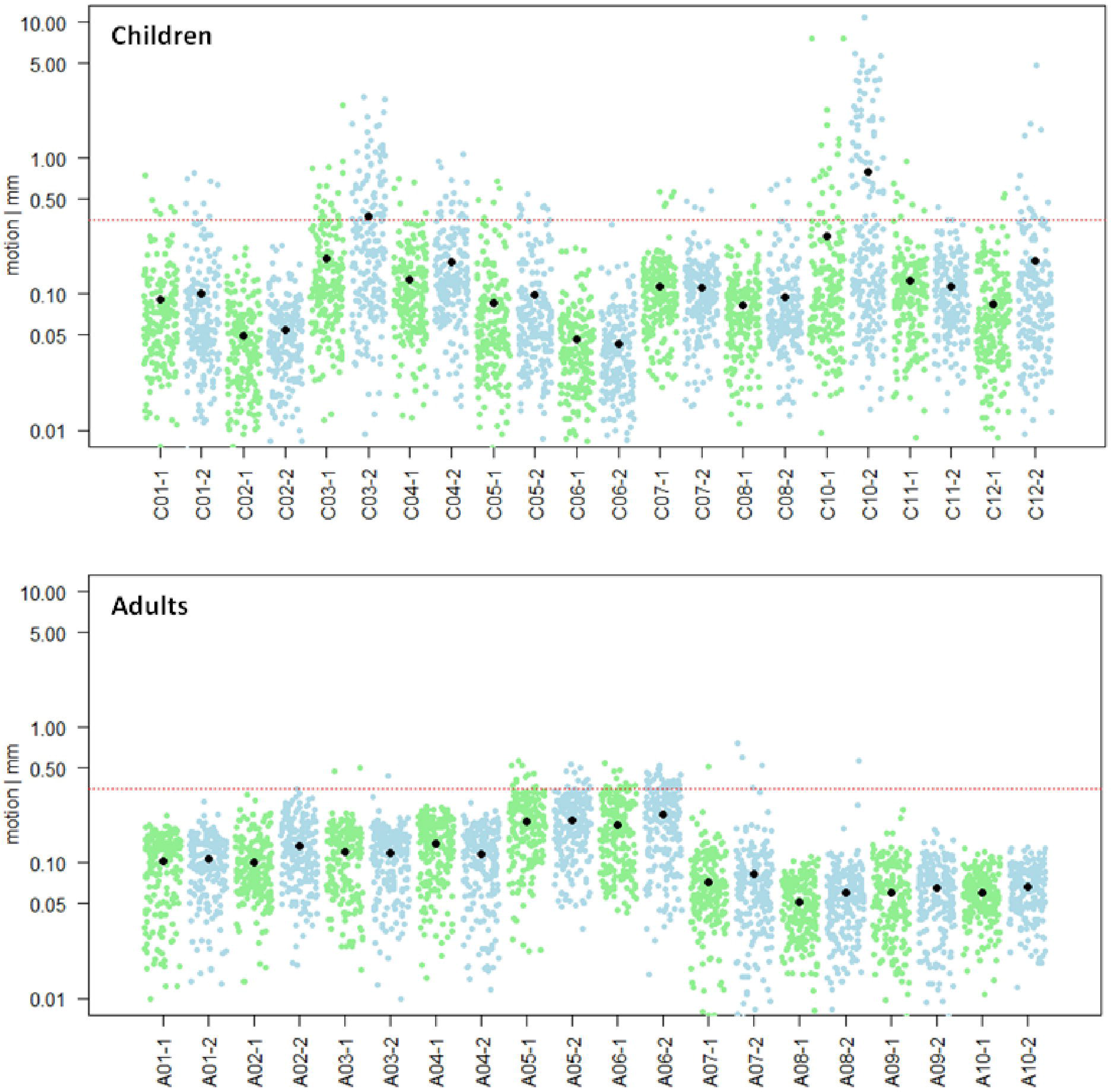
Motion analysis. Mean scan-to-scan motion of the child (n=11, top) and the adult (n=10, bottom) sample, separately for each subject and each session. The first session is depicted in green, the second session in blue. On the x-axis, the subject and session are specified; on the y-axis, the mean scan-to-scan motion is given in logarithmic representation to show normal distribution of data points. The red line marks the cut-off score of a mean scan-to-scan motion threshold of 0.35 mm. The black dots represent the mean scan-to-scan motion. Child C03 and Child C10 were excluded from further analyses due to motion exceeding the predefined session-specific mean scan-to-scan motion threshold of > 0.35 mm in the second session.

### 3.2 Do children activate the core system of face processing?

Our first aim was to assess whether it is possible to detect brain activity in the core system of face processing in children at the single subject level. For illustrational purposes, a representative brain activation pattern is shown in Fig. 4.

**Figure 4:**
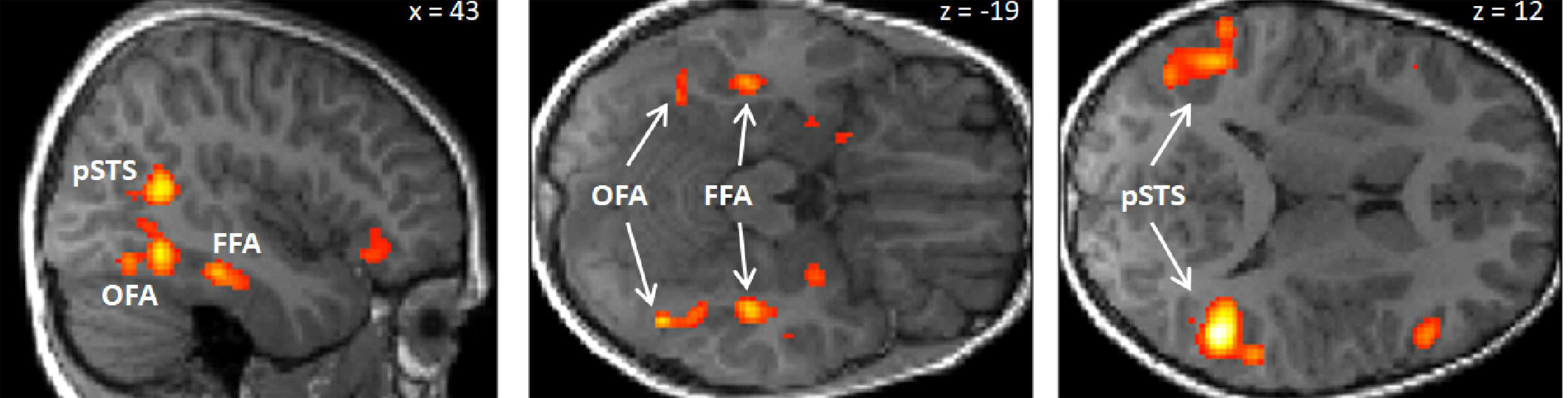
Activation pattern of a representative child (8 years, male) for the contrast “faces > houses”. The subject shows clearly discernible activity in bilateral OFA, FFA and pSTS. For illustrational purposes, the activation pattern is thresholded at p < 0.001, uncorrected for multiple comparisons. Note, however, that all six activations were significant for multiple comparisons (p < 0.05 FWE corrected) at the predefined ROI masks for bilateral OFA, FFA and pSTS.

At the group level, we found in the adults group clearly discernible face-sensitive brain activity in bilateral OFA and bilateral pSTS (p < 0.001, uncorrected). In the children group, we found brain activity in the right OFA and bilateral pSTS at p < 0.001, uncorrected. The left OFA was activated at p < 0.05, uncorrected. In contrast, the left and right FFA was not activated in both groups, not even at a liberal threshold of p < 0.05, uncorrected (see discussion for an explanation). At the individual subject level, all regions of the core system of face processing could be identified in almost all subjects. In most cases, activity was found even at conservative statistical thresholds, i.e., at p < 0.05, FWE corrected for multiple comparisons at the whole-brain level. These results suggest that the core face processing regions, i.e., bilateral OFA, FFA and pSTS can be portrayed at the single-subject level in children and adults, with 100% ROI identification scores of OFA in both samples and slightly lower ROI identification scores of bilateral FFA and pSTS (see Fig. 5 for details).

**Figure 5:**
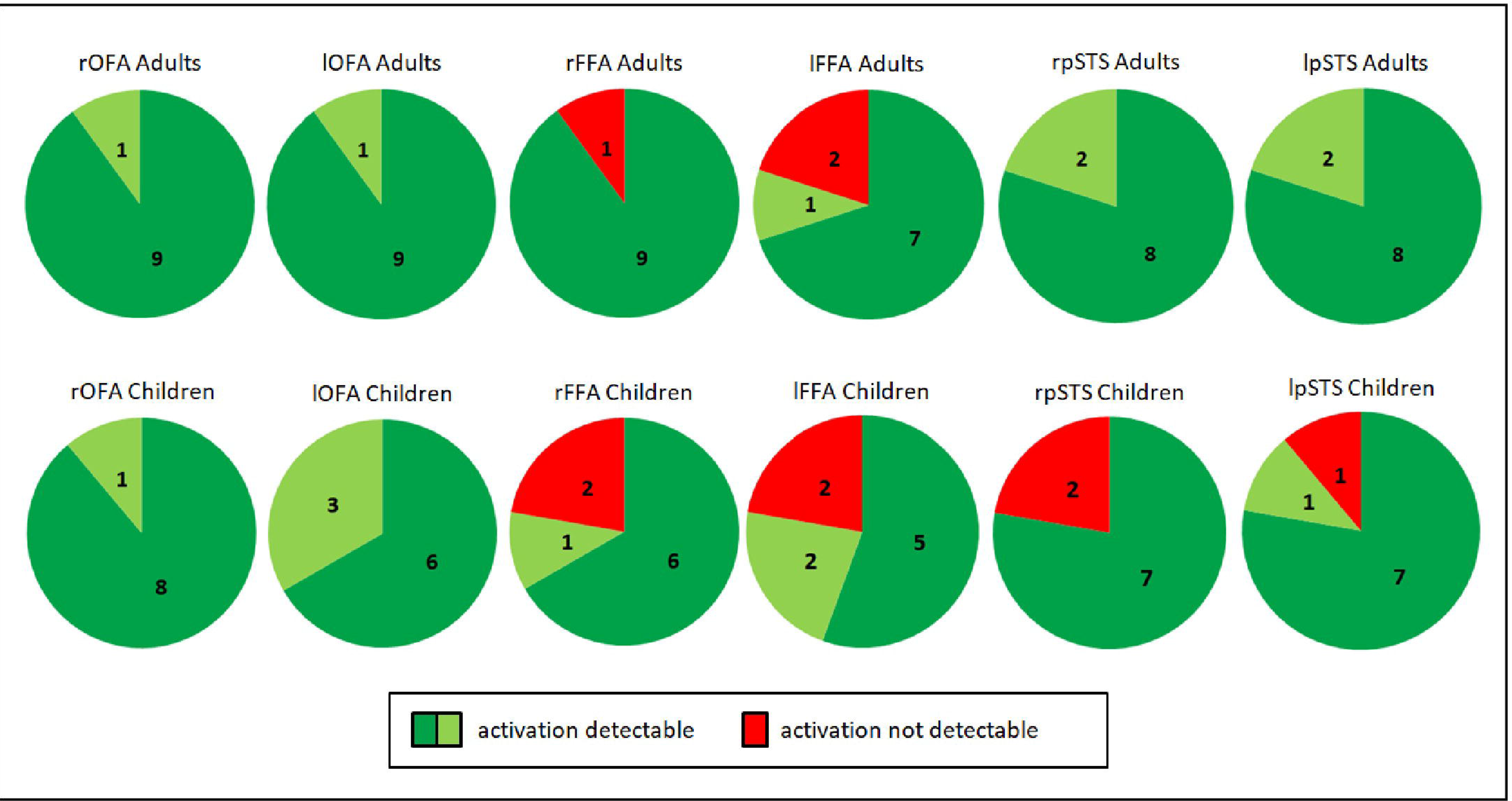
Identification of bilateral OFA, FFA and pSTS in adults (top, n = 10) and children (bottom, n = 9). Dark green indicates activity detectable at a statistical threshold of p < 0.05, FWE corrected for multiple comparisons at the whole brain level. Light green indicates activity detectable at a statistical threshold of p < 0.001 / p < 0.05 uncorrected. Red indicates no detectable activity. In all adults, brain activity could be detected in the right and left OFA as well as in the right and left pSTS. Activity could not be detected in the right FFA for one subject (A02) and the left FFA for two subjects (A02, A07). This was caused by the positioning of the measured volume (see discussion). As for adults, in all children brain activity could be detected in the right and left OFA. Activity was not detected for two children in the right pSTS (C04, C06) and for one child in the left pSTS (C04). Activity was also not detected for two children in the right FFA (C07, C08) and for two children in the left FFA (C01, C08). Missing FFA activity was again caused by the positioning of the measured volume.

### 3.3 Do children and adults differ in brain activation and hemispheric lateralization?

Our second aim was to compare brain activity and hemispheric lateralization in the core system of face processing between adults and children. The mean activation is summarized separately for both groups for the left and right OFA, FFA and pSTS in Fig. 6 (top). Mean activity was, as expected, stronger in bilateral OFA and FFA for adults than for children (albeit the FFA differences were only marginal). In contrast, mean activity for bilateral pSTS was unexpectedly higher for children compared to adults. However, none of the differences reached statistical significance (Table 1). The mean lateralization is summarized in Fig. 6 (bottom). For adults, the mean LI was bilateral for the OFA and right dominant for both FFA and pSTS. For children, the LI was bilateral for both OFA and FFA and right-dominant for the pSTS. Again, none of the differences reached statistical significance (Table 1)2.

**Table 1:**
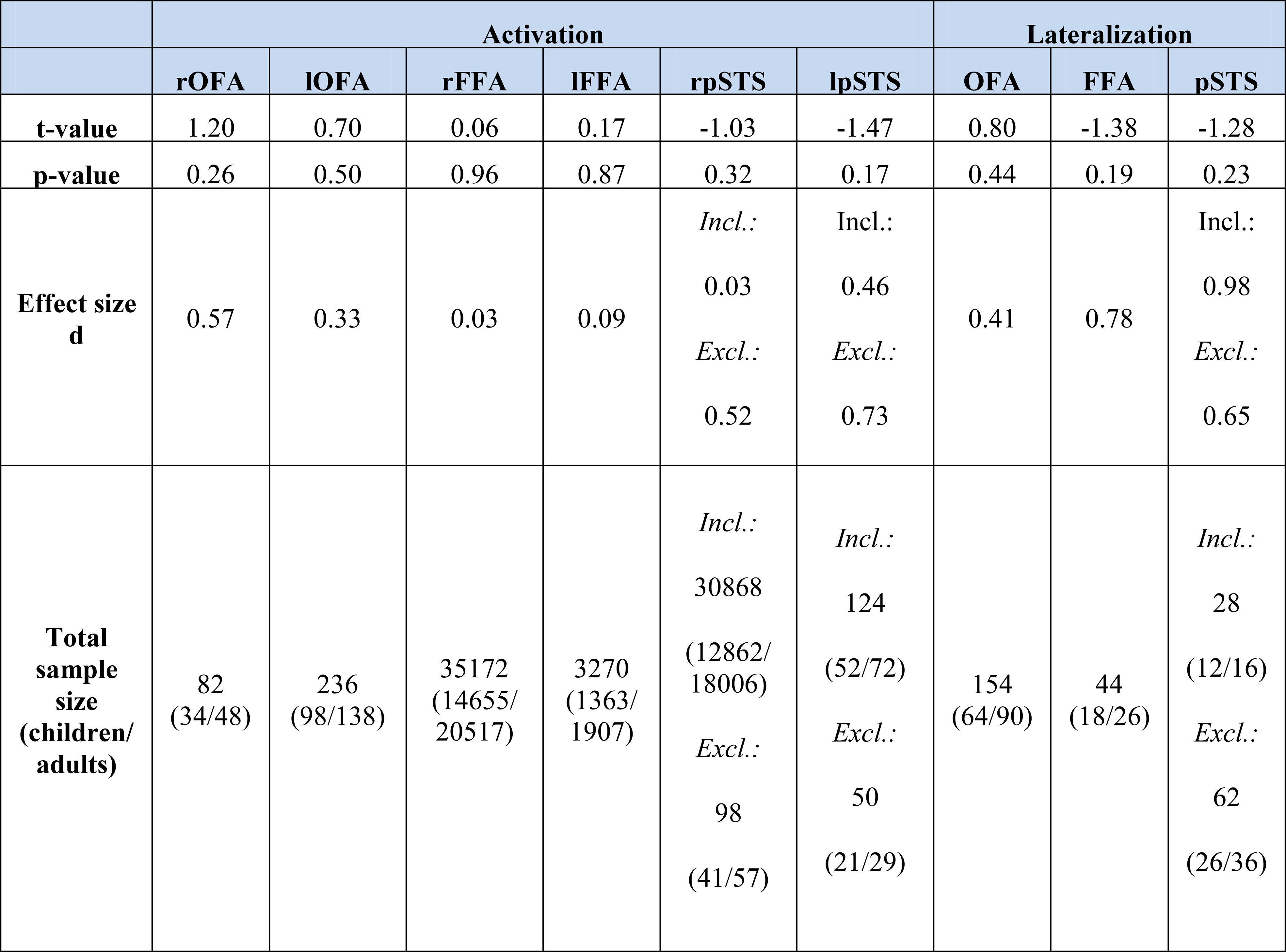
T- and p-values of the Welch t-test, comparing activation and lateralization differences between adults and children for OFA, FFA and pSTS. None of the differences reached statistical significance (p > 0.05). Group differences were used to estimate the effect size Cohen’s d and the sample size for a sufficiently powered follow-up study (statistical power 0.8, alpha error probability 0.05, unbalanced adults/children allocation ratio of 1.4). The effect size and resulting sample size for pSTS activation and lateralization is calculated both inclusive (incl.) and exclusive (excl.) the children without detectable activity (see footnote 3).

**Figure 6:**
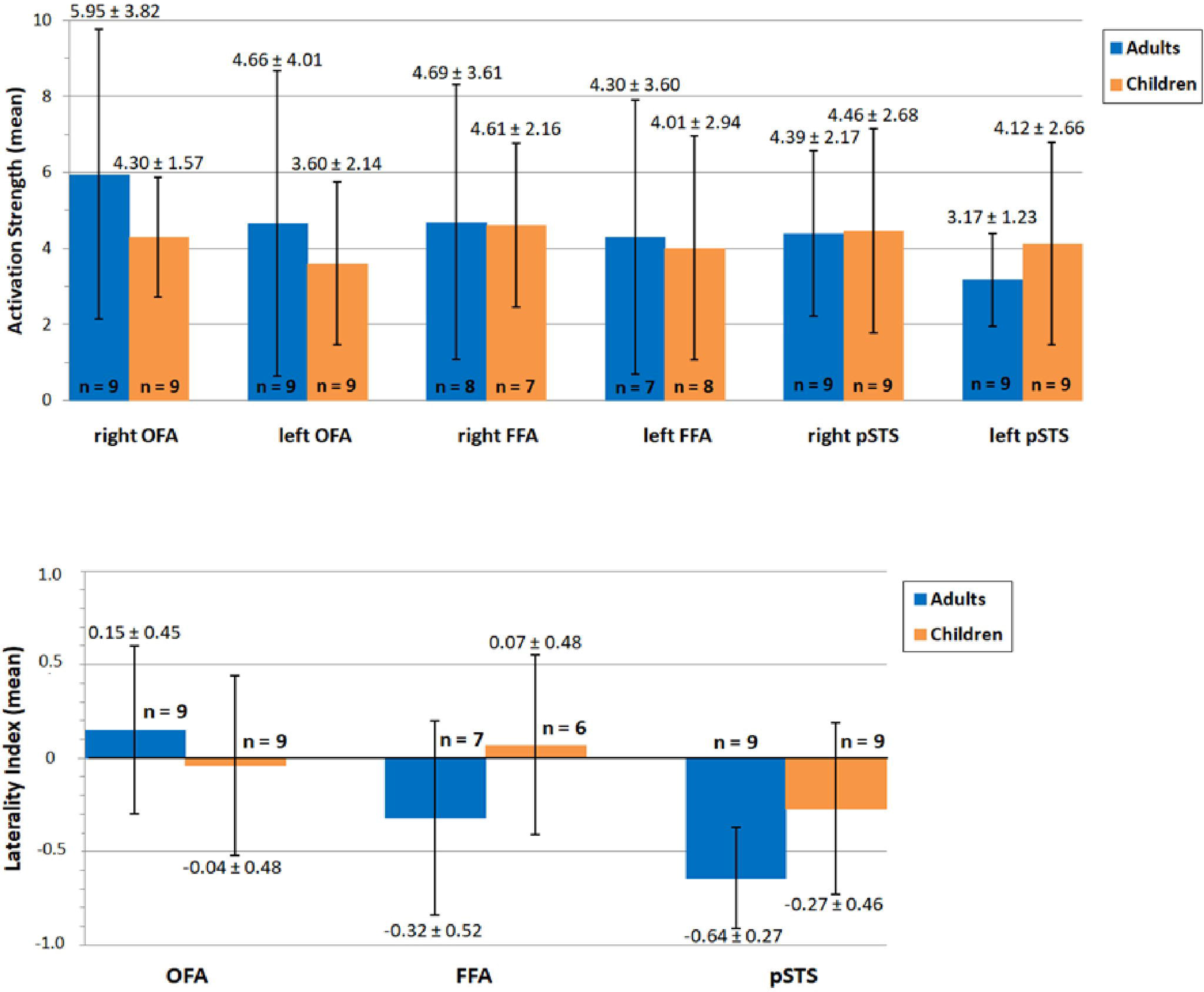
Mean activation (top) and lateralization (bottom) of OFA, FFA and pSTS during face processing in adults (blue) and children (orange). Values of mean activation/ lateralization ± SD are specified above/below error bars. The number of included subjects is depicted at the bottom of/above each bar. None of the differences reached statistical significance (p > 0.05).

The estimated effect size of the activation and lateralization difference between children and adults was assessed by Cohen’s d (Table 1). The activation differences were characterized by effect sizes ranging from 0.03 to 0.57, indicating small to medium effects. In contrast, the estimated lateralization difference between children and adults was characterized by medium to large effect size measures.

The variable of main interest, the hemispheric lateralization of FFA activity, is characterized by an effect size of d = 0.78. Based on these estimates, we performed a power analysis to calculate the group size for a sufficiently powered follow-up study (statistical power 0.8, alpha error probability 0.05, unbalanced adults/children allocation ratio of 1.4). To assess lateralization differences between children and adults in the FFA, the necessary sample size is n = 44 (18 children, 26 adults). The sample size necessary for the assessment of other activation and lateralization differences, respectively, is summarized in Table 1.

## 4 DISCUSSION

In the present study, we established an fMRI paradigm to assess brain activity in the core system of face processing in 7- to 9-year-old children. First, we showed that it is possible to localize the core system’s brain regions in young children even at the single subject level. Second, our results hint at a developmental shift from bilateral FFA activity in children to a right-hemispheric lateralization in adults and thus provide a basis for an informed power analysis for a larger follow-up study. In the following, we will discuss these findings in more detail.

### 4.1 Do children activate the core system of face processing?

Our first aim was to set up a paradigm for assessing activation in the core system of face processing in children at the single subject level. We modified the “standard” fMRI face processing paradigm that we (and others) are using in adults to make it more suitable for the measurement of young children. Much effort was spent on the thorough preparation of the children for the MRI scanning session. The MRI scanning session was put in a child-appropriate frame story. Children slipped into the role of an astronaut on a journey through the universe. Only one child aborted the measurement prematurely due to anxiety, while the other children completed the measurements without problems. When asked after data acquisition, children did not report any feeling of anxiety, but rather curiosity about the device.

Children typically have more difficulties in staying motionless during data acquisition than adults. One of the biggest problems in fMRI of young children are therefore motion artifacts. A recent study from Wilke et al. (2018) reported for instance a positive correlation between proceeding scanning time and the occurrence of motion artifacts in children. We therefore shortened the paradigm, compared to the standard task used in our lab in adults, and additionally split it into two parts with a break of 20 seconds in-between, in order to have the possibility to analyze only data of the first part, if motion artifacts would have increased with proceeding time in the second part. In fact, both parts of the experiment of 9 out of 11 children data could be used thanks to a thorough preparation in advance. The first appointment with the children and their caregivers was crucial, as it gave us the opportunity to convey the importance of lying still to the children. In addition, the few days between the first and second appointment enabled the children to internalize the fMRI procedure. Using a thorough motion analysis, 9 out of 11 children passed the stringent, a-priori chosen motion threshold. The overall motion of the included subjects was comparable between children and adults. This rules out that potential activation differences between children and adults were caused by unequal motion artifacts3.

The left and right OFA was identified in all children and adults. Also the left and right FFA could be localized in most subjects. At first view, it was puzzling that both the left and right FFA were not activated (even at liberal thresholds) at the group level, but in most participants, both adults and children, at the individual subject level even at conservative thresholds. A more detailed analysis showed that this finding could be explained by the positioning of the measured brain volume. As described in the methods section, we did not measure the whole brain, but a “slab” (Fig. 2). The slab was manually positioned with help of the high-resolution structural image. In previous studies of our research group, we used a lateral view of the structural image and oriented the slab at the lowest part of the occipital pole and the lowest part of the inferior temporal gyrus. This positioning ensured that all three regions of the core face processing network (i.e., bilateral OFA, FFA and pSTS) could be measured. In the present study, however, we also aimed, as part of another project, to cover parts of the extended system that are known to play an essential role in emotion processing (in particular the amygdala, insula, cingulate gyrus and inferior frontal gyrus). We therefore used the middle view of the structural image and oriented the slab at the lowest part of the occipital pole and the lowest part of the prefrontal cortex. Due to the different positioning procedure, the FFA was now located at the border of the measured brain volume and was accidentally cropped off in two adults and two children. Since SPM12 does not depict group brain activity in voxels in which at least one subject was not assessed, we were not able to detect activity at the group level. The goal of choosing the defined slab to cover the above-mentioned areas of the extended face processing network in high resolution was a compromise that made it difficult to analyze the data at group level due to “cutting off” the FFA in a few subjects. Future studies therefore have to make sure that all relevant brain regions are included in the measured brain volume. This can be achieved either by measuring a larger brain volume (at the cost of a higher acquisition time per volume (i.e., higher TR) and/or a lower spatial resolution) or choosing a different positioning procedure (orienting the slab at the lowest part of the occipital pole and the lowest part of the inferior temporal gyrus using a lateral view of the structural image). However, if possible, it is advisable to narrow down the target areas and the focus of the study prior to fMRI data acquisition.

pSTS activation was found in all adults, but not in two children. Differently from the FFA, missing activity could not be explained by the positioning of the slab since the pSTS was clearly within the measured brain volume for all subjects. It has been previously reported that it is more difficult to localize the pSTS than OFA and FFA if static stimuli (as in the present study) are used, since the pSTS has been associated with the processing of changeable facial features (e.g., gaze direction, lip movements; Ishai et al., 2005). On the one hand, one might therefore argue that the non-identification of pSTS activity in two subjects is well in line with previous studies using static face localizers (Rhodes et al., 2009; Schultz & Pilz, 2009). On the other hand, it should be noted that we found pSTS activity in the “faces > houses” contrast in all of our adult subjects despite the use of static stimuli. For one of the children subjects that did not exhibit pSTS activity, no activation was found throughout the whole core network during the presentation of faces. This could imply that this child generally processes faces in a different manner compared to the average population. However, the assumption of this child possibly having prosopagnosia is vague, since so far, no corresponding tests have been carried out. It would be favorable for future studies to include a behavioral face recognition test (e.g., Cambridge Face Memory Test) in order to ensure face recognition abilities in general. The pSTS results also indicate that children might apply different face processing strategies than adults, since those children in which we identified pSTS activity showed much higher activation strength in comparison to the adult group (see discussion below).

### 4.2 Do children and adults differ in brain activation and hemispheric lateralization?

Our second aim was to compare brain activation in the core system between children and adults. We hypothesized that children would show reduced activity in all brain regions of the core system (i.e., bilateral OFA, FFA, pSTS) and reduced hemispheric lateralization.

In fMRI, activation differences between two groups are typically assessed by voxel-wise comparisons of normalized functional images. However, in the present study this analysis would have required the use of the same template (e.g., the MNI template) to normalize the data of both children and adults. Since the brains of children and adults largely differ in size and form, this normalization process might have introduced systematic differences between both groups. It would not have been possible to exclude that potential group differences between children and adults both in location and activation strength might simply be related to differences in the normalization process. To surpass this problem, we determined the location of all core system’s brain regions individually for each subject and calculated the activation strength from these regions. Our group comparison thus avoided a potentially biased voxel-wise comparison.

Our results showed, as expected, a trend for weaker activity of children’s bilateral OFA. This finding is in line with theories postulating an increase of face-selective activation in the core system due to an age-related increase of functional specialization (Joseph et al., 2011; Scherf et al., 2007). In contrast, the FFA activity between adults and children was comparable, with an estimated effect size d < 0.1. Unexpectedly, we found for the pSTS a trend for higher activity in children compared to adults (even if we included the children without detectable pSTS activation in the comparison, see results). There are several possible explanations for this hyperactivity. On the one hand, it might be related to the development of the pSTS as a heteromodal association area which builds an interface between sensory signals and hierarchical higher areas (Gogtay et al., 2004). In case of the face-processing network, the bilateral pSTS builds an interface between the core and the extended face-processing network (Hoffman & Haxby, 2000). It might be argued that bilateral pSTS hyperactivation in children is driven through the close link to the hyperactivated extended face-processing network.

On the other hand, pSTS hyperactivation may be, at least in part, explained by change of focus of attention. It can be speculated that the stronger pSTS activity in children is driven by a focus of attention on changeable aspects of the face. Overall, the observed trend for stronger pSTS activation is an interesting finding which deserves further investigation.

The main focus of the present study was the assessment of the development of hemispheric lateralization, in particular of the FFA. Our results showed a trend for increased lateralization of all three regions, with the largest effect found for the FFA. The estimated effect size (d = 0.78) is typically considered to describe medium to large effects, suggesting that differences between children and adults are not negligible. Nonetheless, our findings indicate only a tendency that needs to be underpinned by further data collection with a larger sample size to substantiate the possibility of robustly localizing the core network regions. Using our results as basis for an informed power analysis, we estimated that an adequately powered (sensitivity 0.8) follow-up study testing developmental changes of FFA lateralization would require the inclusion of 18 children and 26 adults. By combining the face processing paradigm with a language task, we are now able to design a follow-up study testing the hypothesis that due to competition between language-biased left-hemispheric VWFA specialization and face representation in the left hemisphere, face representation that was initially bilateral is driven to become right-specialized in the course of development (Behrmann & Plaut, 2015).

Taken together, the results of the present pilot study showed that it is possible to localize the core system in children at the single subject level. They further hint at a developmental shift from bilateral FFA activity in children to a right-hemispheric lateralization in adults. They can be used as basis for an informed power analysis for a larger follow-up study in which we systematically investigate the relationship between the development of reading abilities and hemispheric lateralization of neural activity associated with reading and face processing, respectively.

## Acknowledgments

We are very grateful to Mechthild Wallnig for her help with the MRI data collection.

## Author Contributions

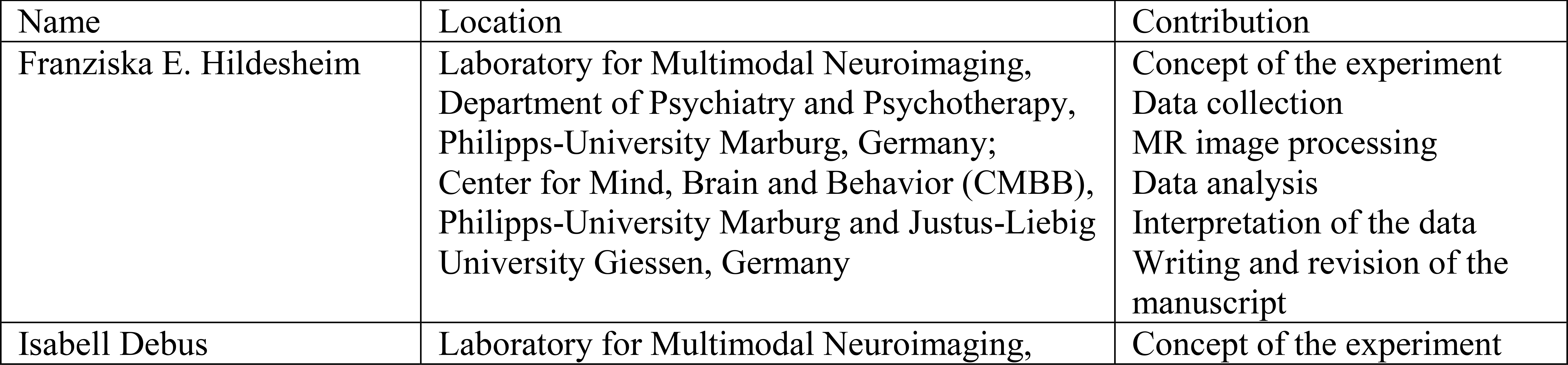
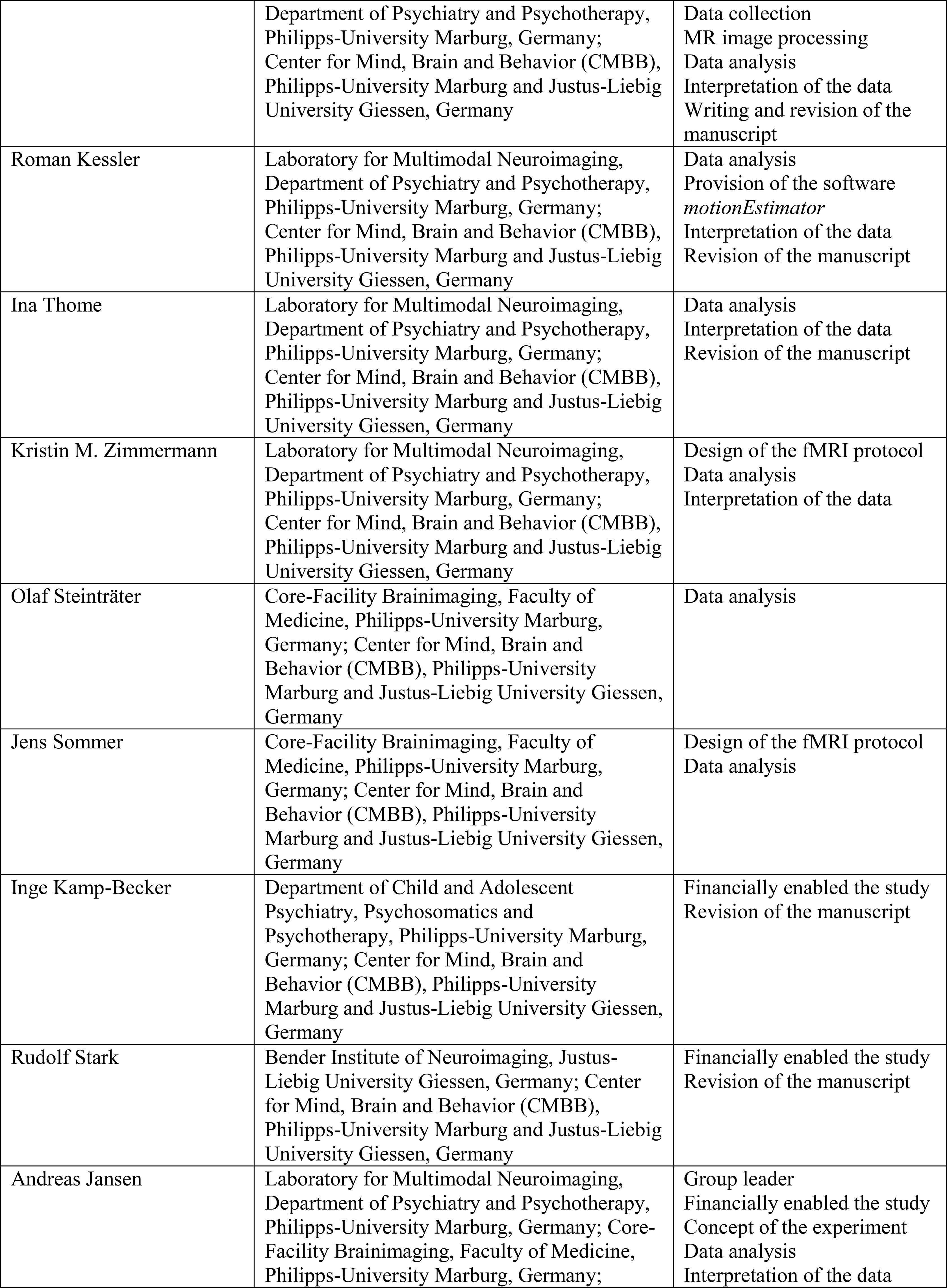
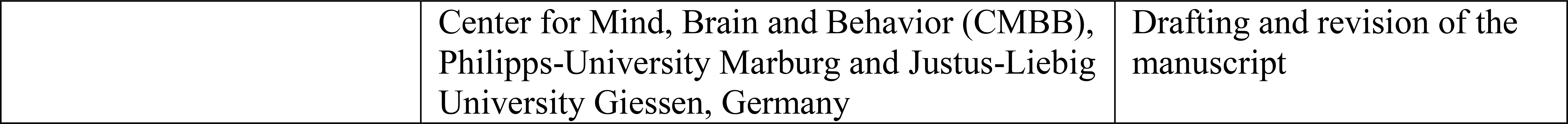

## Footnotes

1 Using this approach, it was of course only possible to calculate brain activity for those subjects in which we were able to find brain activity that was clearly attributable to a core system’s area. We will discuss in the results section the consequences both for the assessment of group differences and the estimation of unbiased effect size measures.

2 At this point we would like to make a methodological remark: pSTS activity could be detected for all adults, but not for two children for the right pSTS and for one child for the left pSTS (Fig. 5). Differently from the FFA, the missing activity could not be explained by the positioning of the slab since the pSTS region was clearly within the measured brain volume for all subjects (see discussion). For a formal comparison of both groups, we therefore had to decide how to deal with this issue. If we decided not to include the two children, the brain activation difference between children and adults would have been potentially overestimated. We therefore decided to also include activity measures for children without detectable pSTS activation. For these subjects, we used the group activation maximum of the right and left pSTS as center for the spherical masks in which we calculated the brain activity. If we had not included the subjects with non-detectable activity in the pSTS, mean activity differences would have been larger (right pSTS: adults 4.39 ± 2.17, children 5.48 ± 2.04; left pSTS: adults 3.17 ± 1.23, children 4.57 ± 2.44). FFA lateralization was only calculated for subjects who had detectable brain activity in both the right and left FFA. For children, pSTS lateralization was calculated on the one hand for all subjects (as before, in case of non-detectable brain activation we used the group activation maximum of the right and left pSTS as center for the spherical masks in which we calculated the brain activity), on the other hand only for those subjects in which brain activity could be found in both the right and left pSTS. The pSTS-LI was −0.27 ± 0.46 if all subjects were included and −0.45 ± 0.31 if only subjects with detectable brain activity were included.

3 We post-hoc analyzed the data of the two excluded children. One child (C03) moved so much that the measured volume was shifted outside the brain regions we were interested in, making it impossible to assess activity in the core system of face processing. The other child (C10) did not show any activity in the core system, also at liberal thresholds (p=0.05 uncorrected).

4 The fMRI paradigm used in the present study was a slightly modified version of the standard paradigm used in our group (e.g., Frässle *et al*., 2016b) to adapt for the assessment of children. First, to minimize the total scanning time, the number of blocks was reduced from 44 to 32. Second, the stimulus presentation time was tripled to 900 ms. Third, the number of stimuli per block was reduced from 20 to 11. Fourth, the number of different face identities was reduced from 30 to 20 (10 female and 10 male identities). The number of different houses was also reduced from 30 to 20, to match it with the number of face stimuli.

## REFERENCES

Anderson, S.F., Kelley, K. & Maxwell, S.E. (2017). Sample-size planning for more accurate statistical power: A method adjusting sample effect sizes for publication bias and uncertainty. Psychological Science, 28(11):1547–1562.

Aylward, E.H., Park, J.E., Field, K.M., Parsons, A.C., Richards, T.L., Cramer, S.C., Meltzoff, A.N. (2005). Brain activation during face perception: Evidence of a developmental change. Journal of Cognitive Neuroscience, 17:308–319.

Baron-Cohen, S., Wheelwright, S., Skinner, R., Martin, J., Clubley, E. (2001). The Autism-Spectrum Quotient (AQ): Evidence from Asperger Syndrome/High-Functioning Autism, Males and Females, Scientists and Mathematicians. Journal of Autism and Developmental Disorders, 31(1):5–17.

Baron-Cohen, S. & Wheelwright, S. (2004). The Empathy Quotient: An Investigation of Adults with Asperger Syndrome or High Functioning Autism, and Normal Sex Differences. Journal of Autism and Developmental Disorders, 34(2):163–175.

Bayer HealthCare (Hg). (2008). Paula in der Röhre. Ein Malbuch für Kinder zur MRT Untersuchung.

Behrmann, M. & Plaut, D.C. (2015). A vision of graded hemispheric specialization. Annals of the New York Academy of Sciences, 1359(1):30–46.

Bernstein, M. & Yovel, G. (2015). Two neural pathways of face processing: A critical evaluation of current models. Neuroscience & Biobehavioral Reviews, 55:536–46.

Bukowski, H., Dricot, L., Hanseeuw, B., Rossion, B. (2013). Cerebral lateralization of face-sensitive areas in left-handers: Only the FFA does not get it right. Cortex, 49(9):2583–2589.

Buxton, R.B., Wong, E.C. & Frank, L.R. (1998). Dynamics of blood flow and oxygenation changes during brain activation: the balloon model. Magnetic Resonance in Medicine, 39(6):855–864.

Cantlon, J.F., Pinel, P., Dehaene, S., Pelphrey, K.A. (2011). Cortical representations of symbols, objects, and faces are pruned back during early childhood. Cerebral Cortex, 21:191–199.

Cohen Kadosh, K., Cohen Kadosh, R., Dick, F., Johnson, M.H. (2011). Developmental changes in effective connectivity in the emerging core face network, Cerebral Cortex, 21(6):1389–1394.

Dehaene, S. & Cohen, L. (2011). The unique role of the visual word form area in reading. Trends in Cognitive Sciences, 15(6):254–262.

Dien, J. (2009). A tale of two recognition systems: Implications of the fusiform face area and the visual word form area for lateralized object recognition models. Neuropsychologia, 47:1–16.

Dima, D., Stephan, K.E., Roiser, J.P., Friston, K.J., Frangou, S. (2011). Effective connectivity during processing of facial affect: Evidence for multiple parallel pathways. Journal of Neuroscience, 31(40):14378–14385.

Dundas, E.M., Plaut, D.C. & Behrmann, M. (2013). The joint development of hemispheric lateralization for words and faces. Journal of Experimental Psychology: General, 142(2):348–358.

Esteban, O., Birman, D., Schaer, M., Koyejo, O.O., Poldrack, R.A., Gorgolewski, K.J. (2017) MRIQC: Predicting quality in manual MRI assessment protocols using no-reference image quality measures. PLoS One.

Fairhall, S.L. & Ishai, A. (2007). Effective connectivity within the distributed cortical network for face perception, Cerebral Cortex, 17(10):2400–2406.

Faul, F., Erdfelder, E., Buchner, A., Lang, A.G. (2009). Statistical power analyses using G*Power 3.1: Tests for correlation and regression analyses. Behavior Research Methods, 41:1149–1160.

Fox, C.J., Iaria, G. & Barton, J.J. (2009). Defining the face processing network: Optimization of the functional localizer in fMRI. Human Brain Mapping, 30:1637–1651.

Frässle, S., Krach, S., Paulus, F.M., Jansen, A. (2016a). Handedness is related to neural mechanisms underlying hemispheric lateralization of face processing. Scientific Reports, 6:1–17.

Frässle, S., Paulus, F.M., Krach, S., Jansen, A. (2016b). Test-retest reliability of effective connectivity in the face perception network. Human Brain Mapping, 37(2):730–744.

Frässle, S., Paulus, F.M., Krach, S., Schweinberger, S.R., Stephan, K.E., Jansen, A. (2016c). Mechanisms of hemispheric lateralization: Asymmetric interhemispheric recruitment in the face perception network. NeuroImage, 124:977–988.

Friston, K.J., Harrison, L. & Penny, W. (2003). Dynamic causal modelling. NeuoImage, 19:1273–1302.

Gathers, A.D., Bhatt, R., Corbly, C.R., Farley, A.B., Joseph, J.E. (2004). Developmental shifts in cortical loci for face and object recognition. Neuroreport, 15:1549–1553.

Gogtay, N., Giedd, J.N., Lusk, L., Hayashi, K.M., Greenstein, D., …, Thompson, P.M. (2004). Dynamic mapping of human cortical development during childhood through early adulthood. PNAS, 101(21):8174–8179.

Golarai, G., Ghahremani, D.G., Whitfield-Gabrieli, S., Reiss, A., Eberhardt, J.L., …, Grill-Spector, K. (2007). Differential development of high-level visual cortex correlates with category-specific recognition memory. NatureNeuroscience, 10:512–522.

Gschwind, M., Pourtois, G., Schwartz, S., Ville, D. van de, Vuilleumier, P. (2012). White-matter connectivity between face-responsive regions in the human brain. Cerebral Cortex, 22(7):1564–1576.

Haist, F., Adamo, M., Wazny, J.H., Lee, K., Stiles, J. (2013). The functional architecture for face processing expertise: FMRI evidence of the developmental trajectory of the core and the extended face systems. Neuropsychologia 51:2893–2908.

Haxby, J.V., Hoffman, E.A. & Gobbini, M.I. (2000). The distributed human neural system for face perception. Trends in Cognitive Science, 4(6):223–233.

He, W., Garrido, M.I., Sowman, P.F., Brock, J., Johnson, B.W. (2015). Development of effective connectivity in the core network for face perception. Human Brain Mapping, 36:2161–2173.

Hemond, C.C., Kanwisher, N.G. & Op de Beeck, H.P. (2007). A preference for contralateral stimuli in human object- and face-selective cortex. PLoS One, 2(6):e574.

Herrington, J.D., Taylor, J.M., Grupe, D.W., Curby, K.M., Schultz, R.T. (2011). Bidirectional communication between amygdala and fusiform gyrus during facial recognition. NeuroImage, 56(4):2348–2355.

Hillger, L.A. & Koenig, O. (1991). Separable mechanisms in face processing: evidence from hemispheric specialization. Journal of Cognitive Neuroscience, 3(1):42–58.

Hoffman, E.A. & Haxby, J.V. (2000). Distinct representations of eye gaze and identity in the distributed human neural system for face perception. NatureNeuroscience, 3(1):80–84.

Ishai, A. (2008). Let’s face it: it’s a cortical network. NeuroImage, 40(2):415–419.

Ishai, A., Schmidt, C.F. & Boesiger, P. (2005). Face perception is mediated by a distributed cortical network. Brain Research Bulletin, 67:87–93.

Jansen, A., Menke, R., Sommer, J., Förster, A.F., Bruchmann, S., Hempleman, J., Weber, B., Knecht, S. (2006). The assessment of hemispheric lateralization in functional MRI – Robustness and reproducibility. NeuroImage, 33(1):204–217.

Joseph, J.E., Gathers, A.D. & Bhatt, R.S. (2011). Progressive and regressive developmental changes in neural substrates for face processing: Testing specific predictions of the interactive specialization account. Developmental Science, 14:227–241.

Jospe, K., Flöel, A. & Lavidor, M. (2018). The interaction between embodiment and empathy in facial expression recognition. Social cognitive and affective neuroscience, 13(2):203–215.

Li, J., Liu, J., Liang, J., Zhang, H., Zhao, J., …, Lee, K. (2010). Effective connectivities of cortical regions for top-down face processing: A Dynamic Causal Modeling study. Brain Research, 1340:40–51.

Lundqvist, D., Flykt, A. & Öhman, A. (1998). The Karolinska Directed Emotional Faces – KDEF, CD ROM from Department of Clinical Neuroscience, Psychology section, Karolinska Institutet, ISBN 91-630-7164-9.

Maldjian, J.A., Laurienti, P.J., Kraft, R.A., Burdette, J.H. (2003). An automated method for neuroanatomic and cytoarchitectonic atlas-based interrogation of fMRI data sets. NeuroImage, 19(3):1233–1239.

Mandeville, J.B., Marota, J.J., Ayata, C., Zaharchuk, G., Moskowitz, M.A., …, Weisskoff, R.M. (1999). Evidence of a cerebrovascular postarteriole windkessel with delayed compliance. Journal of Cerebral Blood Flow & Metabolism, 19(6):679–689.

Mechelli, A., Price, C.J., Friston, K.J., Ishai, A. (2004). Where bottom-up meets top-down: Neuronal interactions during perception and imagery. Cerebral Cortex, 14(11):1256–1265.

Meng, M., Cherian, T., Singal, G., Sinha, P. (2012). Lateralization of face processing in the human brain, Proceedings of the Royal Society: Biological Sciences, 279(1735):2052–2061.

Morawetz, C., Holz, P., Lange, C., Baudewig, J., Weniger, G., Irle, E., Dechent, P. (2008). Improved functional mapping of the human amygdala using a standard functional magnetic resonance imaging sequence with simple modifications. Magnetic Resonance Imaging, 26(1):45–53.

Müller, V.I., Cieslik, E.C., Turetsky, B.I., Eickhoff, S.B. (2012). Crossmodal interactions in audiovisual emotion processing. NeuroImage, 60(1):553–561.

Nagy, K., Greenlee, M.W. & Kovács, G. (2012). The lateral occipital cortex in the face perception network: An effective connectivity study. Frontal Psychology, 3:141.

Naumann, S., Senftleben, U., Santhosh, M., McPartland, J., Webb, S.J. (2018). Neurophysiological correlates of holistic face processing in adolescents with and without autism spectrum disorder. Journal of Neurodevelopmental Disorders, 10(1):27.

Nguyen, V.T., Breakspear, M. & Cunnington, R. (2014). Fusing concurrent EEG-fMRI with dynamic causal modeling: Application to effective connectivity during face perception. NeuroImage, 102:60–70.

Oldfield, R.C. (1971). The assessment and analysis of handedness: The Edinburgh Inventory. Neuropsychologia, 9(1):97–113.

Passarotti, A.M., Paul, B.M., Bussiere, J.R., Buxton, R.B., Wong, E.C., Stiles, J. (2003). The development of face and location processing: an fMRI study. Developmental Science, 6(1):100–117.

Peelen, M.V., Glaser, B., Vuilleumier, P., Eliez, S. (2009). Differential development of selectivity for faces and bodies in the fusiform gyrus. Developmental Science, 12:F16–F25.

Power, J.D., Schlaggar, B.L. & Petersen, S.E. (2015). Recent progress and outstanding issues in motion correction in resting state fMRI. NeuroImage, 0:536–551

Price, C.J. & Devlin, J.T. (2003). The myth of the visual word form area. NeuroImage, 19:473–481.

Rhodes, G., Brake, S. & Atkinson, A.P. (1993). What’s lost in inverted faces? Cognition, 47(1):25–57.

Rhodes, G., Michie, P.T., Hughes, M.E., Byatt, G. (2009). The fusiform face area and occipital face area show sensitivity to spatial relations in faces. European Journal of Neuroscience, 30:721–733.

Rossion, B. (2015) Face perception. In: Toga, A.W. (edt.). Brain Mapping: An Encyclopedic Reference, 2nd edn., pp. 515–522. Academic Press: Elsevier.

Scherf, K.S., Behrmann, M., Humphreys, K., Luna, B. (2007). Visual category-selectivity for faces, places and objects emerges along different developmental trajectories. Developmental Science, 10:F15–F30.

Schultz, J. & Pilz, K.S. (2009). Natural facial motion enhances cortical responses to faces. Experimental Brain Research, 194:465–475.

Schuster, V., Herholz, P., Zimmermann, K.M., Westermann, S., Frässle, S., Jansen, A. (2017). Comparison of fMRI paradigms assessing visuospatial processing: Robustness and reproducibility. PLoS One, 12(10):e0186344.

Schwarzer, G. (2000). Development of face processing: The effect of face inversion. Child Development, 71:391–401.

Song, Y., Zhu, Q., Li, J., Wang, X., Liu, J. (2015). Typical and atypical development of functional connectivity in the face network, Journal of Neuroscience, 35(43):14624–14635.

Wechsler, D. & Naglieri, J.A. (2006). Wechsler Nonverbal Scale of Ability. San Antonio, TX: Harcourt Assessment.

Wilke, M. & Schmithorst, V.J. (2006). A combined bootstrap / histogram analysis approach for computing a lateralization index from neuroimaging data, NeuroImage, 33(2):522–530.

Wilke, M. & Lidzba, K. (2007). LI-tool: A new toolbox to assess lateralization in functional MR-data, Journal of Neuroscience Methods, 163(1):128–136.

Wilke, M., Groeschel, S., Lorenzen, A., Rona, S., Schuhmann, M.U. (2018). Clinical application of advanced MR methods in children: points to consider. Annals of Clinical and Translational Neurology, 5(11):1434–1455.

Willems, R.M., Peelen, M.V. & Hagoort, P. (2010). Cerebral lateralization of face-selective and body-selective visual areas depends on handedness. Cerebral Cortex, 20(7):1719–1725.

Yovel, G., Tambini, A. & Brandman, T. (2008). The asymmetry of the fusiform face area is a stable individual characteristic that underlies the left-visual-field superiority for faces. Neuropsychologia, 46:3061–3068.

